# DeepBranchAI: A Novel Cascade Workflow Enabling Accessible 3D Branching Network Segmentation

**DOI:** 10.64898/2026.03.25.714249

**Authors:** Alexander V. Maltsev, Lisa M. Hartnell, Luigi Ferrucci

**Affiliations:** Intramural Research Program, National Institute on Aging, Baltimore, MD, United States

**Keywords:** cascade training, 3D deep learning, annotation efficiency, topological segmentation, transfer learning, branching networks, random forests, nnU-Net

## Abstract

Three-dimensional branching networks exist throughout biological, natural, and man-made systems as pathways through volumetric space. Segmentation is required to correctly reconstruct the networks in whole or in part for analysis. This presents a unique challenge as minor voxel misclassifications can cause sporadic connectivity shifts, whereby connected elements appear to disconnect (false negatives) or to even become amplified (false positives). Addressing this topological vulnerability requires the generation of 3D models since 2D slice-by-slice approaches cannot maintain connectivity across x, y, and z axes. Yet tracking 3D architecture demands substantially more analytical resources than using a 2D strategy as generating volumetric annotations requires extraordinary amounts of expert time to manually annotate. This creates a fundamental annotation bottleneck: with sparse training data available, deep learning models tend to overfit available volumes and fail to generalize to novel volumes. We present a cascade training workflow that overcomes this bottleneck through a positive feedback loop in which trained models become annotation aids for subsequent volumes. The workflow begins with random forests that generate initial drafts from minimal labels, followed by expert refinement that cycle ever closer to the ground truth. As refined data accumulates, training transitions from 2D to 3D architectures, which systematically expand sparse datasets into comprehensive training sets. The outcome is a 3D nnU-Net model optimized for topology-preserving segmentation. We dub our resulting model DeepBranchAI. Training validation on heavily branching mitochondrial networks, generated by focused ion beam scanning electron microscopy (FIB-SEM, 15nm voxel resolution) achieved Dice Similarity Coefficient (DSC) = 0.942 across 5-fold cross-validation. Transfer learning to vascular networks (VESSEL12 dataset, CT volumes, 30,000-fold voxel size difference) achieved 97.05% accuracy against ground truth, validating that learned features represent domain-general topological principles. This workflow reduces annotation time from months to weeks while transforming sparse initial labels into robust training sets. Complete implementation, trained weights, and validation code are provided open source.

## 1. Introduction

Mapping the connectivity of 3D branching networks is fundamental to understanding their function and integrity, yet achieving accurate segmentation remains prohibitively difficult. These structures appear throughout biological, natural and engineered systems, such as mitochondria and vasculature in tissues, porous membranes, 3D-printed lattices in materials, river networks and root systems. Whether naturally occurring or engineered, these networks typically solve a universal challenge: to efficiently distribute resources throughout space while minimizing costs to the system. The essential first step for measuring relationships in any such system is to determine which components connect to which others and how these relationships are arranged in space. Volumetric imaging requires segmentation, identifying which voxels belong to network structures versus background. Expert hand-traced annotation is the most reliable mapping method of connectomes, but processing complex volumes can consume over 50 person-years of manual proofreading [1, 2]. Convolutional neural networks (CNN) present sophisticated computer-automated support as a solution to this bottleneck.

However, training CNNs requires large amounts of accurately labeled data and producing that data takes the very expert time the automation is supposed to conserve. Branching networks are also unusually difficult to segment due to their topological fragility. Small errors in identifying which voxels belong to the network can significantly alter perceived network architecture, including the number of branches, connectivity patterns, and network topology, leading to flawed conclusions about system properties [3]. This topological vulnerability necessitates 3D segmentation models. Only with volumetric context can one properly capture and measure three-dimensional connectivity patterns.

The problem is that volumetric annotation takes orders of magnitude longer than 2D labeling. With limited training data, deep learning models overfit and memorize what they’ve seen and fail on new volumes. Existing approaches typically apply 2D methods slice-by-slice and sacrifice topological accuracy or invest heavily into annotation [4, 5]. The central question is whether this bottleneck for 3D model training and annotation can be overcome through well-designed AI workflow design rather than through ever-increasing manual labor.

Our approach combines conventional machine learning (ML) with deep learning (DL), using each where it’s strongest. When training data is sparse, conventional machine learning algorithms use standard image features, such as random forests and support vector machines, to achieve reasonable accuracy [6, 7]. Deep learning algorithms require substantial initial training data but scale effectively, yielding progressively higher accuracy through the modeling of complex, hierarchical image features. Our cascade approach uses conventional machine learning to rapidly bootstrap preliminary annotations, and then through expert feedback iteratively refines these annotations to generate robust ground truth for training deep neural networks. Once trained, the deep learning models produce better drafts than the conventional ML, and each further round of annotation becomes easier. The models become annotation assistants. This dramatically reduces annotation requirement while maintaining quality, making the creation of large-scale 3D training datasets substantially more efficient.

We imaged mitochondrial networks using focused ion beam scanning electron microscopy (FIB-SEM), which provides nanometer-scale resolution (15 nm isotropic voxels) acceptable to capture structural details of interest to us [8, 9]. Initially, we developed this workflow for use in segmenting mitochondrial networks in skeletal muscle tissue. We found that they exhibit all canonical difficulties in 3D network segmentation: thin interconnections easily broken apart or falsely amplified by voxel misclassification, variable contrast across imaging volumes, complex branch points, and topological vulnerability [10, 11, 12]. These challenges are present in many 3D branching systems [10, 11, 12], making our developed method broadly transferable to applications in materials science, geophysics, neuroscience, and engineering.

Using this workflow, we produced our neural net model DeepBranchAI, a 3D nnU-Net model optimized for segmenting branching networks [13]. We validated that our model is generalizable by successfully applying transfer learning to segment vascular networks in CT volumes, which have similar topological branching but through different imaging techniques (CT X-ray attenuation vs. FIBSEM electron scattering), different spatial scales (30,000-fold voxel volume difference), and describes a different biological system.

## 2. Related Work

Deep learning has achieved remarkable success in volumetric segmentation, building on foundational architectures like U-Net [4] and its 3D extensions [5], with specialized models such as DeepVesselNet [11] targeting vascular networks and TotalSegmentator [14] enabling comprehensive CT/MRI organ segmentation. TotalSegmentator is trained on the 3D nn-UNet architecture just as our DeepBranchAI. However, 3D models need substantially more training data than their 2D counterparts and creating high-quality annotated training data demands extraordinary investments of expert time. With only sparse training data available, deep learning models overfit to available samples and fail to generalize to novel volumes. Recent approaches address this challenge through pseudo-label generation, semi-supervised learning, and iterative bootstrapping strategies that expand sparse annotations into dense segmentations [15, 16]. However, these methods typically remain within deep learning frameworks and aim to minimize human involvement entirely.

Methods like clDice loss [17] and connectivity-constrained objectives [3] have improved segmentation of branching structures, but they don’t help with the annotation problem because they still need extensive, topology-consistent ground truth to begin with. Transfer learning [18, 19] reduces annotation requirements but typically assumes domain similarity. Our work, on the other hand, explicitly tests transfer across extreme domain boundaries, including 30,000-fold differences in scale and distinct imaging modalities. Our cascade training framework takes a complementary approach: rather than improving model learning efficiency, it improves annotation generation efficiency through strategically designed human–AI workflows. Unlike purely automated bootstrapping methods that try to cut human involvement out, our framework explicitly transitions from conventional machine learning to deep learning and treats expert refinement as a feature rather than a limitation. The resulting workflow generalizes across domains where topology-sensitive networks require faithful 3D reconstruction (Table 1).

**Table 1:** Domain Applications of Topology-Preserving 3D Segmentation.

| Domain | Target Structure | Imaging Modality | Critical Topology Challenge |
| --- | --- | --- | --- |
| Materials Science | Porous membranes | Micro-CT | Permeability depends on connected pore paths [20, 21] |
| Geophysics | Fracture networks | Seismic volumes | Connectivity determines fluid flow [22, 23] |
| Neuroscience | Neural circuits | EM volumes | Single voxel errors break synaptic connections [24, 25] |
| Plant Biology | Root systems | MRI/CT | Network connectivity affects nutrient transport [26, 27] |
| Engineering | 3D printed lattices | X-ray CT | Structural integrity requires continuous struts [28, 29] |
| Cell Biology | Mitochondrial networks | FIB-SEM | Energy distribution requires network continuity [30, 31] |
| Vascular Biology | Vascular networks | CT | Blood flow requires topological accuracy [32, 33] |

Table 1. Domain applications of topology-preserving 3D segmentation. Each field uses distinct imaging modalities and different target structures but share the critical requirement that segmentation errors must not break 3D connectivity. This makes them ideal candidates for topology-enforcing cascade workflows.

## 3. Methods

### 3.1 Overview of the Cascade Training Framework

Our approach for segmenting 3D branching networks combines three main stages (Figure 1). Stage A (Preprocessing and Training Set Curation) prepares a heterogeneous training set from FIB-SEM image volumes. Stage B (Training Set Generation) iteratively builds and refines high-quality ground truth through a cascade approach, beginning with conventional machine learning for preliminary segmentation, followed by expert refinement cycles that transition to deep learning. Stage C (Training the DeepBranchAI Classifier) employs 3D nnU-Net configurations optimized for topology preservation.

**Figure 1.**
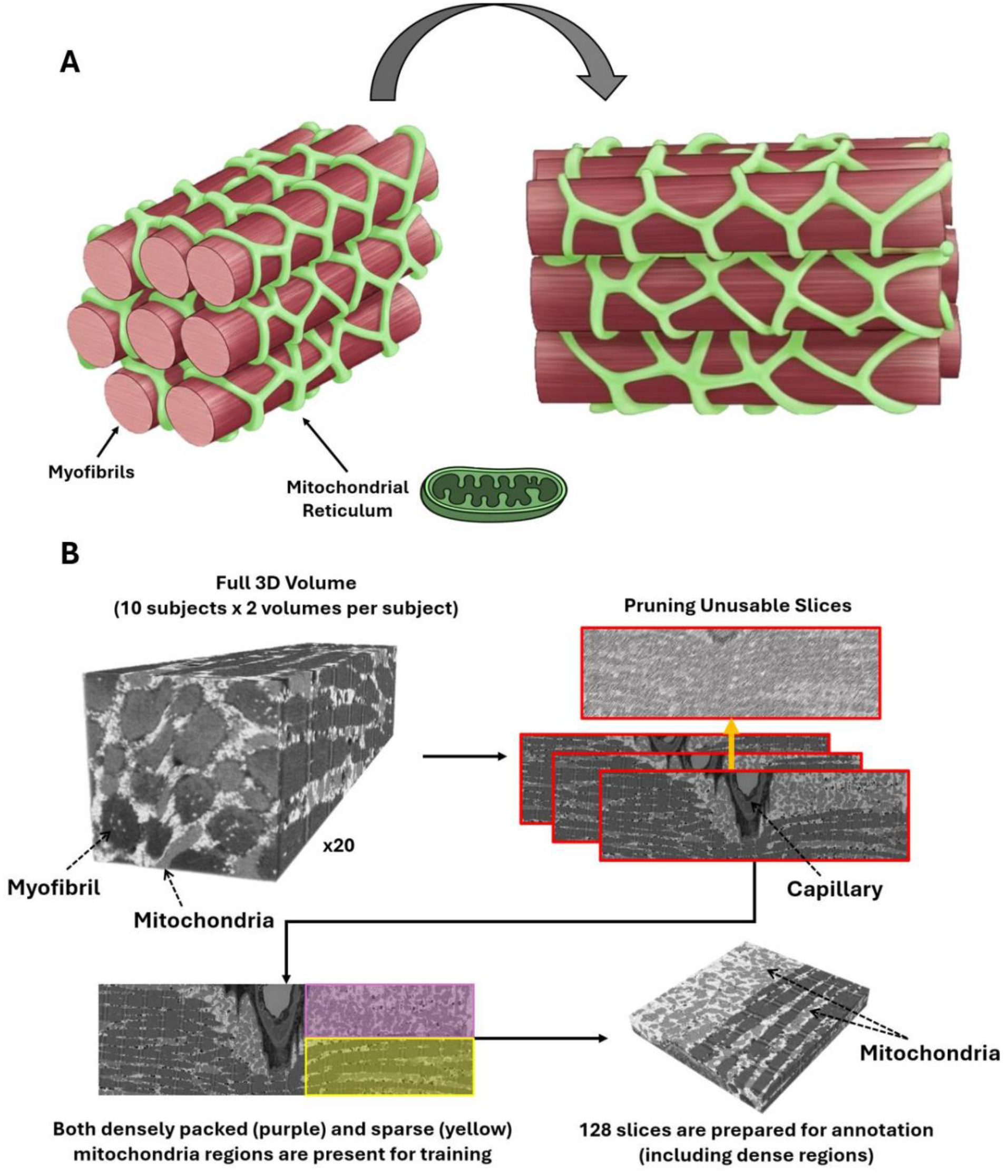
Cascade Training Framework Stage 1: Volume Preparation. (A) Schematic showing mitochondria interspersed between myofibrils in human skeletal muscle. Many of them are in contact with each other, forming a nearly continuous branching network. (B) Preprocessing includes cross-correlation alignment to ensure 3D continuity and artifact removal to prevent spurious training signals. Optional frequency-domain denoising improves signal-to-noise ratio (SNR) while preserving edges. The training set was curated to capture topological diversity. It is visually evident that densely packed mitochondria near capillaries (purple) are organized differently from regions with more linear, sparse arrangements (yellow). Recognizing this diversity allows for sufficient depth of 3D context learning, and more balanced representation to prevent class bias.

### 3.2 Software and Environment

Segmentation was performed using the following software environment:

- Python: 3.12.2
- PyTorch: 2.2.1 with CUDA 11.8
- nnU-Net: v2.3.1
- ORS Dragonfly: 4.0 (for manual annotation and refinement, https://dragonfly.comet.tech/)
- ImageJ/Fiji: with Trainable Weka Segmentation plugin (for initial 2D segmentation)
- Python packages: nibabel, tifffile, numpy, multiprocessing
- Destriping: aind-smartspim-destripe package (https://github.com/AllenNeuralDynamics/aind-smartspim-destripe)

### 3.3 Preprocessing and Training Set Curation

We built the training set to be deliberately heterogeneous. It includes image volumes from processed samples of skeletal muscle biopsies from 10 participants. Images were obtained as TIFF stacks using FIB-SEM, then aligned and binned to produce volumes with 15 nm isotropic resolution. [8, 34]. Both images and labels were converted to NIfTI format (.nii.gz) using the Python *nibabel* library, with segmentation labels stored as binary masks.

#### 3.3.1 Preprocessing Steps

Raw stacks undergo three preprocessing operations (Figure 1):

1. **Cross-correlation and volume alignment** corrects spatial misalignments between slices, ensuring structures remain continuous throughout the volume rather than falsely fragmented.
2. **Interference slice removal** identifies and removes erroneous TIFF slices that disrupt continuity through automated detection of intensity anomalies or manual inspection.
3. **Denoising** optionally applies wavelet-FFT filtering to remove stripe artifacts common in scanning microscopy [35]. The filter uses a Daubechies-3 wavelet with two parameter configurations: cellular images (σ=64, max_threshold=3) and non-cellular regions (σ=128, max_threshold=12). Implementation is provided in Destriping.ipynb.

#### 3.3.2 Training Set Balancing

This curation phase creates a balanced dataset focusing on continuous 3D branching structures (Figure 1):

4. **Sub-structure verification** ensures that features are correctly identified through expert visual inspection. For branching networks, this means including different topological configurations (clustered vs. distributed patterns, peripheral vs. central locations).
5. **Depth requirement** establishes a minimum threshold of 128 Z-slices (∼10% of the maximum z depth of our volume). This gave minimum sufficient volumetric context for learning 3D topology (Figure 1A). Adjust this threshold based on feature of interest structure size and imaging resolution.
6. **Class balancing** provides approximately equal representation of structural sub-types by cropping more training patches from underrepresented regions.

### 3.4 Training Set Generation Through Cascade Workflow

The cascade approach leverages conventional machine learning and deep learning in complementary roles. Conventional ML algorithms achieve reasonable accuracy with limited training data by training with globally effective pre-defined image features, while deep learning requires more initial input but achieves superior accuracy with scale using convoluted features [6, 19].

#### 3.4.1 Initial Conventional ML Stage

7. **Preliminary segmentation** uses the 2D Weka Trainable Segmentation algorithm [36] (ImageJ implementation). Training regions are marked by drawing on representative target and background. This step requires minimal input (typically 5-10 minutes of annotation) and uses conventional ML’s strength to generalize from sparse labels.
8. **Expert refinement** corrects obvious errors through manual editing using ORS Dragonfly 4.0. Corrections are saved as overlays and converted to binary masks. Refinement and segmentation should be done in three dimensions because different axis can reveal different branching morphology.
9. **Iterative retraining** repeats steps 7-8 until reaching saturation, determined through expert validation when visual inspection shows no obvious errors remaining. Typically 2-3 iterations suffice.
10. **High-quality extraction** identifies the most accurate segments through expert visual inspection. The goal is to identify at least 32 consecutive XY slices with high-quality annotations to begin 2D deep learning training.

#### 3.4.2 Transition to Deep Learning

11. **2D neural network training** uses the extracted segments to train a 2D nnU-Net model, marking the transition from conventional ML to deep learning.
12. **Inference on full dataset** applies the trained 2D network to generate probability maps across the entire dataset. These maps guide experts to focus on ambiguous regions.
13. **Final expert refinement** thresholds probability maps to produce initial binary predictions, which experts can further correct to create ground truth for the 3D neural network (typically chosen as 0.50 [13]).

### 3.5 Training DeepBranchAI Classifier

In stage 3 the 3D nnU-Net is trained using a chunk size of 352 × 352 ×128 voxels. We termed the resulting model:DeepBranchAI (Figure 3). This chunk size captures both broad contextual information and fine-grained features necessary for topology preservation.

**Figure 2.**
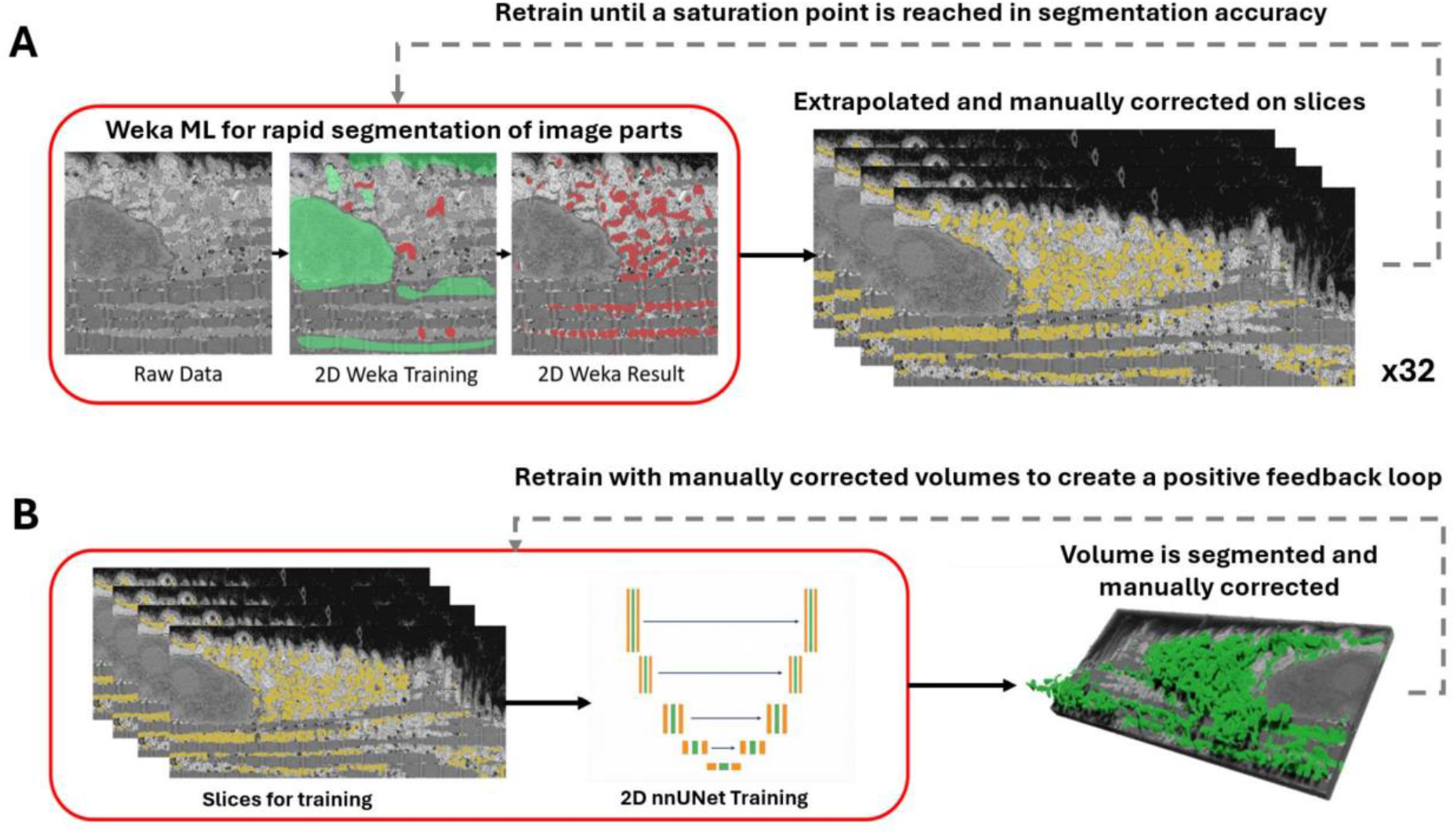
Cascade Training Framework Stage 2: Iterative Ground Truth Construction. (A) The process starts with a 2D Weka random forest classifier trained on minimal annotations. Experts correct the output and retrain iteratively until accuracy plateaus. (B) The refined segments are then used to train a 2D nnU-Net, which generates probability maps across full volumes. Experts do a final refinement pass on these maps, producing ground truth volumes that feed back into further training.

**Figure 3.**
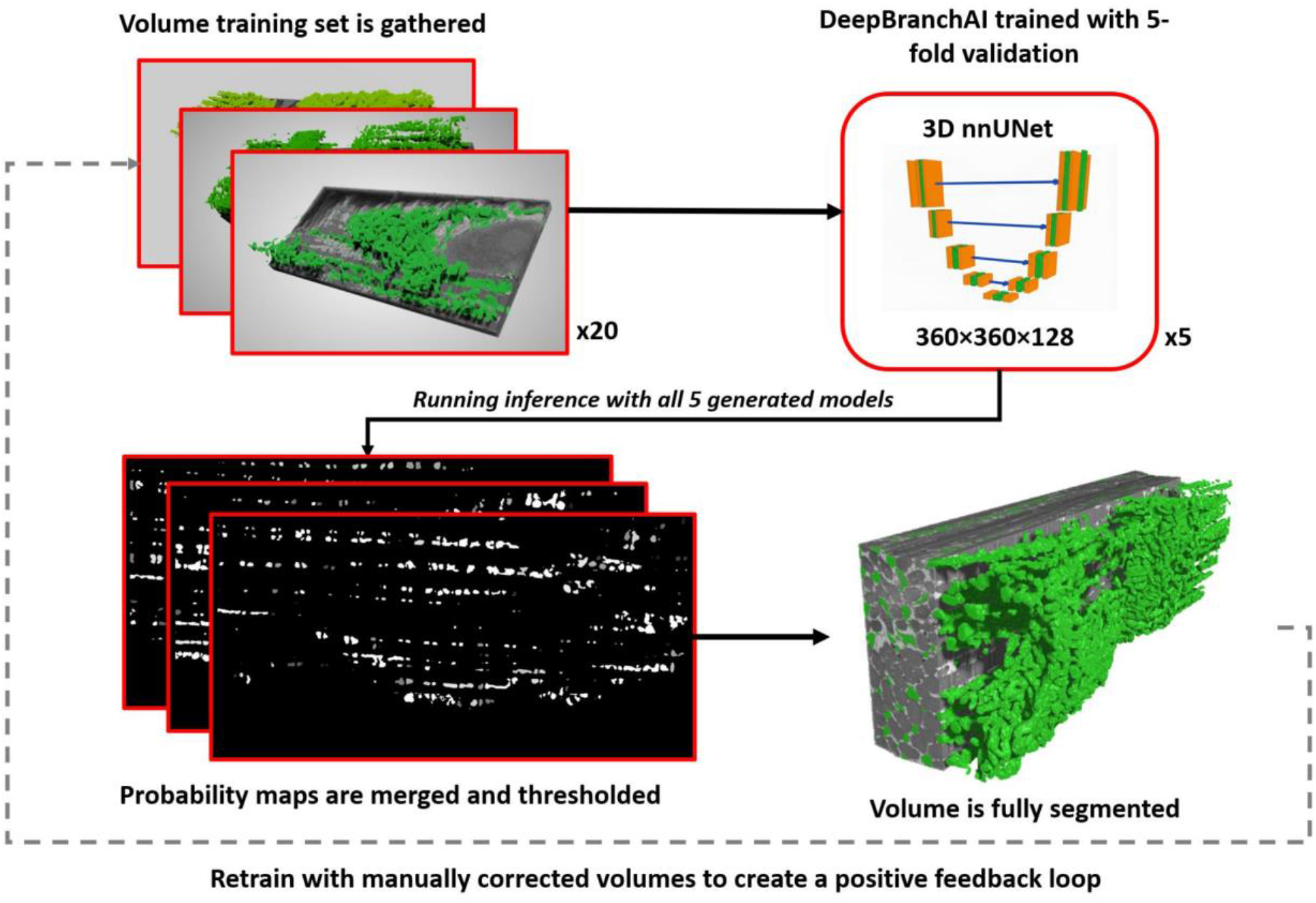
Cascade Training Framework Stage 3: DeepBranchAI and Deployment. The 3D nnU-Net model that we dub DeepBranchAI is trained using high-quality ground truth s with chunk sizes large enough to capture local details and long-range topological relationships. Trained models segment new volumes via chunk-wise processing, generating probability maps that are stitched and thresholded to produce final segmentations.

#### 3.5.1 Training and Inference

14. **3D neural net training** uses the automatic hyperparameter configuration from nnU-Net for 100 epochs with 5-fold cross validation to evenly split our ground truth from 20 volumes into validation hold-out groups of size 4. The chunk size should be scaled proportionally with available GPU memory: 48 GB VRAM accommodates 352 × 352 ×128 voxels, while 24 GB VRAM accommodates approximately 50% of this volume. The nnU-Net algorithms automatically adapt to available memory.
15. **Automated inference** applies all 5 trained models from 5-fold cross-validation via the inference algorithms from nnU-Net, which handles chunk-wise processing with overlapping 3D patches. The final segmentation volume is assembled via stitching and weighted averaging.
16. **Thresholding** applies a 0.50 cutoff to the merged probability map to generate the final binary segmentation.
17. **Optional refinement** applies connected component analysis (26-connected neighborhood) and minimum size thresholding to remove small spurious detections. Additional manual refinement may be performed with the option to retrain the model, creating the positive feedback loop described in the workflow.

### 3.6 Code Availability and Implementation Resources

The complete codebase is provided through Jupyter notebooks:

#### Data Preprocessing

- Demo_Destripe.ipynb: Wavelet-FFT filtering for stripe artifact removal with parallelized processing.
- Training Data Preprocessing.ipynb: Quality control including dimension verification and intensity statistics.

#### Model Training and Evaluation

- nnUNET 3D Train.ipynb: End-to-end training for 3D nnU-Net models
- nnUNET 3D Inference.ipynb: Batch inference for generating 3D segmentation predictions.
- Validate 3D DeepBranchAI.ipynb: Evaluation metrics for 3D models with volumetric analysis.

#### Transfer Learning Validation

- Finetune_VESSEL12.ipynb: Complete pipeline for transfer learning validation including annotation loading, upscaling, and accuracy computation.

All notebooks and trained model weights are available at https://github.com/alexmaltsev/DeepBranchAI.

### 3.7 Evaluation Metrics

We employed six standard metrics to assess segmentation performance [16, 37]. Let TP, TN, FP, and FN denote true positives, true negatives, false positives, and false negatives, respectively:

**Sensitivity (Recall)** measures the proportion of actual target pixels correctly identified:

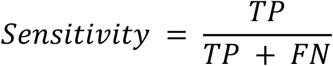

**Specificity** measures the proportion of background pixels correctly identified:

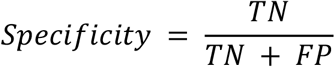

**Accuracy** measures overall proportion of correctly classified pixels:

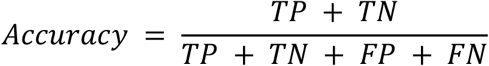

**Dice Similarity Coefficient (DSC)** measures overlap between predicted and ground truth segmentations [17]:

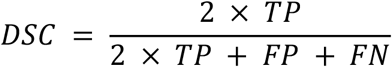

**Absolute Volume Difference (AVD)** quantifies relative difference in total segmented volume:

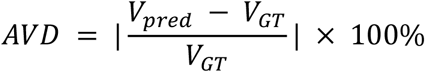

where *V*_*pred*_ and *V*_*gt*_ represent the predicted and ground truth volumes, respectively.

**Cohen’s Kappa (κ)** measures agreement while accounting for chance:

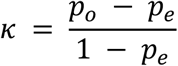

where *p*_*o*_ is observed agreement and *p*_*e*_ is expected agreement by chance. Values above 0.8 indicate excellent agreement.

### 3.8 Computational Requirements

All procedures were performed on a workstation with an NVIDIA RTX A6000 (48 GB VRAM), 128 GB RAM, and 24 logical processors running Ubuntu 24. Typical training times: approximately 30 minutes per iteration for 2D Weka segmentation, 4-6 hours for 2D nnU-Net training, and 2-3 days for complete 3D nnU-Net training on the full dataset of 20 volumes.

### 3.9 Troubleshooting

- **Out-of-memory errors**: nnU-Net automatically reduces patch size. Ensure PyTorch is installed with CUDA support matching your GPU driver.
- **Holes or fragments in segmentation**.: Adjust minimum size threshold in connected component analysis based on expected structure dimensions.
- **Low accuracy scores**: Check class balance in your training data. If one class dominates, the model will learn to predict it by default. If balance is already present, annotate a few more volumes before retraining. **Segmentation accuracy varies across volumes**: The training data doesn’t cover the full range of contrast in your dataset. Make sure destriping is applied to all volumes.
- **Visual inspection does not confirm accuracy**: Dice and IoU can miss topological errors. Try Cohen’s Kappa for as a fallback option or a topology-sensitive metric like dlDice, which penalizes connectivity breaks directly. **Software compatibility**: Ensure correct installation of Python 3.12.2, PyTorch 2.2.1, and nnU-Net v2.3.1. Configure environment variables (nnUNet_raw, nnUNet_preprocessed, nnUNet_results) before training.

## 4. Results

We validate the cascade training framework in two ways. First, we evaluate performance quantitatively across FIB-SEM volumes with different imaging conditions. Second, we test whether the model transfers across domains, which would suggest it has learned something general about 3D branching topology rather than just memorizing features specific to our data.

### 4.1 Quantitative Evaluation of DeepBranchAI

DeepBranchAI demonstrated strong performance in segmenting complex branching networks from FIB-SEM volumetric data with folds ranging from DSC = 0.93 to 0.96 (Table 2)

**Table 2:** 5-fold cross-validation performance for DeepBranchAI (3D nnU-Net)

| Fold | Sensitivity | Specificity | Accuracy | DSC | AVD (%) | $\kappa$ |
| --- | --- | --- | --- | --- | --- | --- |
| 1 | 0.964 | 0.994 | 0.991 | 0.956 | 2.63 | 0.951 |
| 2 | 0.862 | 0.999 | 0.989 | 0.920 | 12.65 | 0.914 |
| 3 | 0.972 | 0.995 | 0.993 | 0.962 | 1.93 | 0.958 |
| 4 | 0.880 | 0.996 | 0.982 | 0.920 | 8.87 | 0.910 |
| 5 | 0.946 | 0.994 | 0.988 | 0.951 | 4.10 | 0.944 |
| Average | 0.925 | 0.996 | 0.989 | 0.942 | 6.04 | 0.935 |

#### 4.1.1 Cross-Validation Performance

We used 5-fold cross-validation with an 80/20 train-validation split, grouping volumes by subject so that no subject appeared in both sets within a fold. The narrow range of DSC scores (0.920-0.962) and consistently high specificity (>0.994) indicates stable performance independent of specific data splits (Table 2). Average sensitivity was 0.925 and specificity 0.996, meaning the model misses relatively little and rarely hallucinates structure. The average Cohen’s kappa of 0.935 indicates excellent agreement between predicted and expert segmentations.

#### 4.1.2 Comparison with 2D and 3D Architectures

Below we compare DeepBranchAI against 2D and 3D U-Net [2, 18] and 2D nnU-Net (Table 3) to check whether full volumetric processing is worth the overhead.

**Table 3:** Performance comparison of different segmentation approaches.

| Model | Sensitivity | Specificity | Accuracy | DSC | AVD | $\kappa$ |
| --- | --- | --- | --- | --- | --- | --- |
| 2D U-Net | 0.721 | 0.971 | 0.947 | 0.726 | 9.26 | 0.697 |
| 3D U-Net | 0.856 | 0.993 | 0.978 | 0.888 | 8.86 | 0.875 |
| 2D nnU-Net | 0.880 | 0.986 | 0.976 | 0.879 | 8.87 | 0.865 |
| DeepBranchAI (3D nnU-Net) | 0.925 | 0.996 | 0.989 | 0.942 | 6.04 | 0.935 |

DeepBranchAI scored highest across all metrics (Table 3). The 2D U-Net achieves the lowest DSC of 0.726, logically because to its inability to capture volumetric context. The 3D U-Net shows much improved performance (DSC 0.888), demonstrating the importance of 3D information for branching structure segmentation [18]. The 2D nnU-Net improves over standard 2D U-Net (DSC 0.879 vs 0.726) through automatic hyperparameter optimization, but still falls short of 3D performance. DeepBranchAI achieves the highest performance and lowest average volume difference (6.04% compared to 8.86-9.26% for other methods), driven by to its ability to capture both in-plane details and volumetric context.

### 4.2 Validation of Cross-Domain Generalizability

To test whether learned features represent fundamental branching topology rather than domain-specific texture patterns, we tested transfer across maximum domain distance: different imaging techniques (CT X-ray attenuation vs. FIBSEM electron scattering), different spatial scales (30,000-fold voxel volume difference), and different biological systems (vascular vs. organelle networks).

The VESSEL12 dataset [10], comprising 23 CT volumes of vascular networks, was used for validation. Ground truth segmentation masks were generated using TotalSegmentator [15] with the “lung vessels” task, refined by applying lung-specific masks to isolate the lung region. Both CT volumes and segmentation masks were upscaled by a factor of 2 using cubic interpolation to approximately match the volumetric pixel dimensions of our FIB-SEM training data.

#### 4.2.1 Transfer Learning Strategy

We employed transfer learning [19, 30] using the pretrained DeepBranchAI model, continuing training from pretrained weights with all parameters allowed to adapt. Training proceeded for 100 epochs using 35% of the total VESSEL12 dataset, strategically selected to include image chunks containing at least 2.0% vessel labels. This subset represents the most vessel-dense regions, ensuring that training was concentrated on information-rich areas rather than predominantly empty background.

#### 4.2.2 Validation Results

This strategy yielded a DSC of 0.88. Furthermore, to provide independent validation beyond TotalSegmentator-derived ground truth, we performed point-by-point validation on expert annotations across three held-out test volumes (VESSEL12 Volume 21, 22, and 23), demonstrating 97.05% overall accuracy (91.81% on vessel pixels, 99.50% on non-vessel pixels).

The successful transfer from mitochondrial networks at nanometer resolution to vascular networks at dramatically different scales validates that the cascade training framework produces models learning generalizable topological principles (branching) rather than domain-specific artifacts (Figure 5).

**Figure 4.**
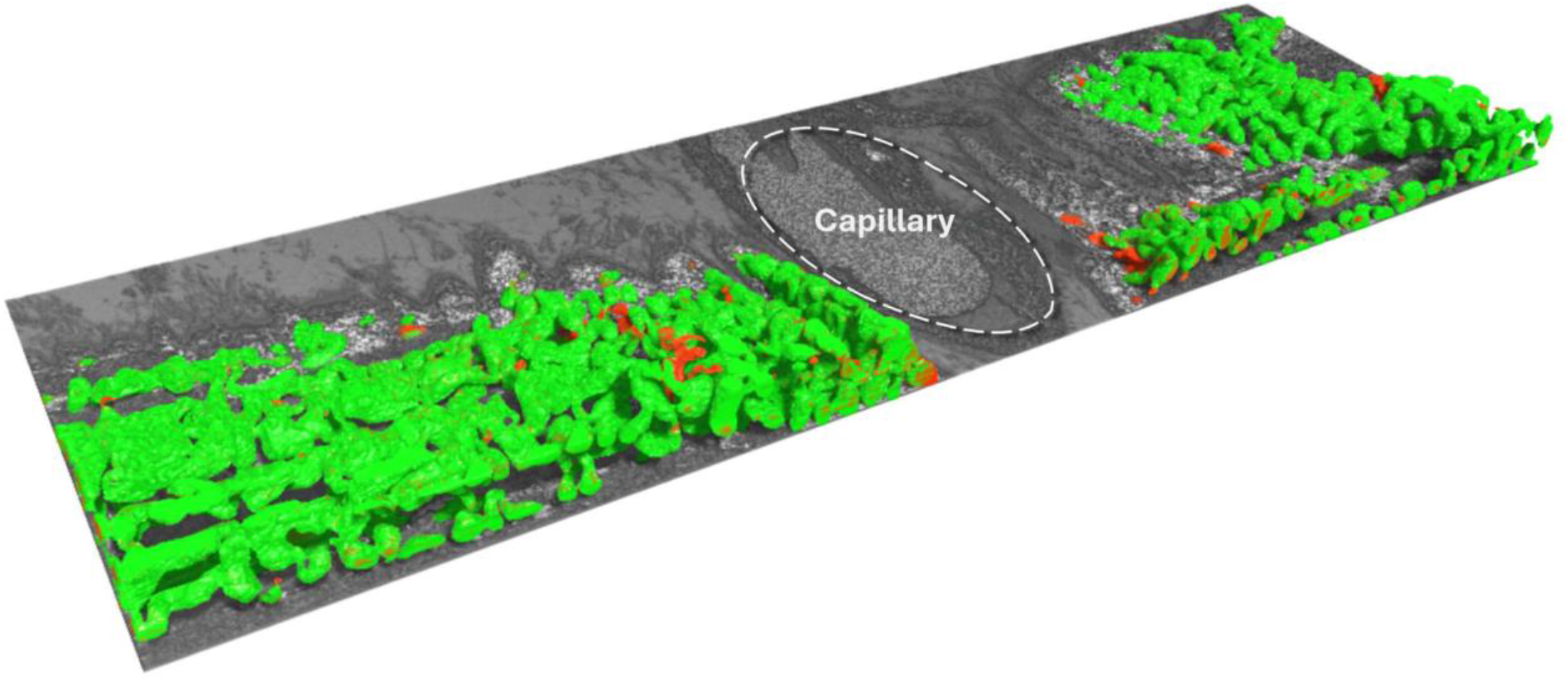
DeepBranchAI Segmentation Performance. Pixel-to-pixel comparison (DSC = 0.97) between DeepBranchAI segmentation (green) and ground truth (red) showing high overlap and discrepancies primarily contained in highly dense regions near the capillary.

**Figure 5.**
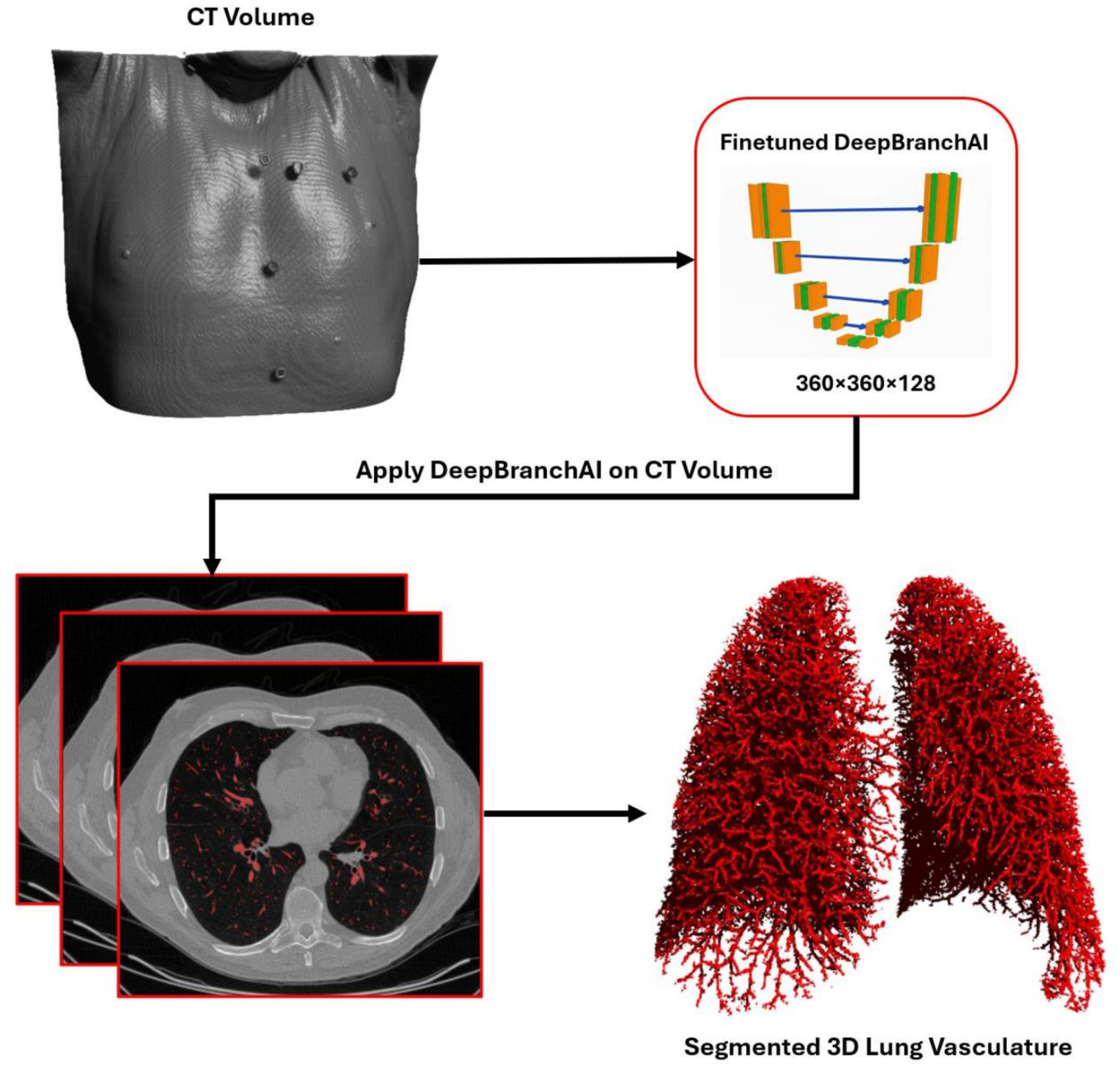
Cross-Domain Transfer Learning Validation on Lung Vasculature. Cross-sectional CT image from VESSEL12 with vascular networks segmented in red. The 3D rendering demonstrates preserved topological connectivity across the entire volume. Fine-tuning on 35% of data achieved 97.05% accuracy against expert annotations.

## 5. Discussion

Segmenting 3D branching networks requires true volumetric models rather than 2D slice-by-slice approaches. Branching structures exhibit topological fragility where even minor voxel misclassifications can break connectivity or amplify structures, and only 3D architectures can recognize that structures appearing disconnected in one slice reconnect in adjacent slices. On the other hand, 3D models need far more training data, and there is no purely automated way around that bottleneck. Our cascade training framework addresses this challenge by combining conventional machine learning, deep learning, and human expertise. Conventional machine learning rapidly generates broad coverage from minimal examples and deep learning achieves high accuracy through learned features.Meanwhile, human experts ensure topological correctness that neither automated approach can guarantee alone. Trained models produce draft annotations superior to conventional ML, creating a positive feedback loop where models become annotation assistants for subsequent volumes. In practice, this cuts annotation time from months to weeks without sacrificing quality.

We validated our approach using mitochondrial networks in skeletal muscle [20, 21]. The 2D U-Net scored 0.726 DSC and the 2D nnU-Net reached 0.879, but neither maintained topological consistency across slices. DeepBranchAI achieved 0.942 DSC, representing a 30% relative improvement, demonstrating its capability for learning spatial 3D relationships essential for recognizing and maintaining network connectivity. Our 352 × 352 ×128 voxel patches span approximately 2 μm in depth, holding the spatial context necessary for this recognition. Our transfer learning results provide additional validation of this approach. Finetuned DeepBranchAI, achieved 97.05% segmentation accuracy against expert annotations for vascular networks in CT volumes from the VESSEL12 dataset [7], despite 30,000-fold differences in voxel volumes, completely different imaging techniques (FIB-SEM versus CT), and different biological systems (mitochondria versus blood vessels). This success aligns with recent findings by Tetteh et al. that show that deep networks trained on tubular vessel structures learn generalizable features rather than domain-specific textures [13]. The extreme scale difference in our validation suggests that this workflow-level consistency produces features capturing fundamental branching geometry that transfer across imaging modalities. These principles likely extend to other branching network types, as the geometric constraints governing resource distribution networks are shared across biological and engineered systems (Table 1).

Our cascade workflow relates to but differs from existing annotation-reduction. Human-in-the-loop methods operate entirely within deep learning frameworks. Cellpose 2.0 demonstrated that iterative correction requires only 100-200 ROIs versus 200,000 for full training [39], MONAI Label provides interactive 3D segmentation with active learning [37], and the AIDE framework achieves comparable results using only 35% of annotations through cross-model self-correction [15]. However, these approaches assume sufficient initial annotations to train a functional neural network. DeepBranchAI addresses this bootstrapping problem by starting with conventional machine learning and transitioning to deep learning only after sufficient ground truth accumulates. While most iterative methods refine segmentation within individual volumes, our framework operates across the dataset, transforming annotation effort from linear to sublinear scaling, making subsequent annotation easier with each new volume.

The positive feedback loop present in this study is evidence to adopt scaling strategies that take advantage of human-AI workflows. However, there are several limitations in this framework. The minimum depth requirement of 128 slices for DeepBranchAI (approximately 2 μm at 15 nm resolution for FIB-SEM) excludes structures below this threshold. Layered or crystalline structures would need a different approach because our transfer learning method works best between networks that specifically share branching topology. Open questions also remain about how to choose the right volumetric context size, and whether metrics like graph edit distance or connectivity preservation at branch points would evaluate topology better than Dice alone. A clear next step is integrating topology-preserving metrics like clDice and cbDice into the workflow. Ultimately, human judgment will remain essential for validating topological correctness in ambiguous cases where automated methods lack ground truth.

## 6. Conclusion

The annotation bottleneck in 3D imaging is solvable, but not by cutting humans out of the loop. Deep learning needs ground truth that it can’t generate on its own. Our framework solves this by starting with conventional ML to bootstrap initial annotations from minimal labels, then transitioning to deep learning as the training set grows through expert-corrected iterations. We validated this framework on mitochondrial network segmentation in skeletal muscle (DSC=0.942) and demonstrated successful transfer to vascular network segmentation in CT volumes (97.05% accuracy using only 35% of target data), confirming that learned representations capture domain-general topological principles rather than dataset-specific features. All code, trained weights, preprocessing pipelines, and documentation are available at https://github.com/alexmaltsev/DeepBranchAI. This workflow should apply wherever 3D connectivity matters, from neural circuit mapping to geological structures to plant phenotyping to materials science. Rather than replacing expert knowledge, effective automation must multiply its impact, and this work provides the tools to implement such workflows.

## 7. Future Developments

Foundation models like SAM [40] and its prompt-based extensions [41] could accelerate initial annotation, but current implementations are fundamentally 2D and cannot enforce the 3D topological consistency that branching networks require. A 3D foundation model trained on diverse volumetric data does not yet exist, and developing one may be the most impactful direction for this field. Collaborative repositories for validated ground truth, analogous to those in genomics and structural biology, would also accelerate progress. Most 3D annotation currently happens in isolation. A platform supporting community contribution, expert validation, and standardized formatting would allow groups to build on each other’s work [2]. Synthetic data generation could also reduce dependence on manual annotation, provided topological properties are preserved. Procedurally generated branching networks with controlled complexity could provide large volumes of topology-consistent training data [11]. Current patch-based training, while necessary for GPU memory constraints, sacrifices global context. Emerging sparse convolution approaches and transformer architectures designed for volumetric data may enable whole-volume training that captures long-range dependencies without prohibitive memory requirements. These directions converge on the same underlying problem of reducing the tradeoff between annotation effort and computational cost that currently constrains 3D segmentation.

## Data Availability

All implementation code, trained model weights, and Jupyter notebooks are available at https://github.com/alexmaltsev/DeepBranchAI under CC0 license. Pre-trained weights are provided in nnU-Net v2.3.1 format for immediate deployment or fine-tuning.

## Acknowledgments

The authors wish to thank the members of the GESTALT biopsy team and participants.

## Author Contributions

AVM: Conceptualization, Methodology, AI Software, Formal analysis, Writing. LMH: Conceptualization, Methodology, FIBSEM Microscopy, Sample Processing, Expert Validation, Writing. LF: Funding acquisition, Project administration, Resources, Supervision, Writing.

## Funding

This research was supported by the Intramural Research Program of the National Institutes of Health (NIH). The contributions of the NIH author(s) are considered Works of the United States Government. The findings and conclusions presented in this paper are those of the author(s) and do not necessarily reflect the views of the NIH or the U.S. Department of Health and Human Services.

## Competing Interests

The authors declare that the research was conducted in the absence of any commercial or financial relationships that could be construed as a potential conflict of interest.

## Abbreviations

DSC: Dice Similarity Coefficient
FIB-SEM: Focused Ion Beam Scanning Electron Microscopy
CNN: Convolutional Neural Network
ML: Machine Learning
nnU-Net: No-new-U-Net (self-configuring neural network framework)
RF: Random Forest
IoU: Intersection over Union
AVD: Absolute Volume Difference

## Key Terms

**Segmentation -** The process of partitioning volumetric image data into regions corresponding to distinct structures or objects of interest.

**Voxel -** A three-dimensional pixel; the smallest distinguishable unit in volumetric imaging data.

**Topological fragility -** The vulnerability of branching networks where minor voxel misclassifications can break connectivity and fragment continuous structures.

**Annotation -** The process of manually labeling structures in images to create training data for machine learning models.

**Cascade training -** A multi-stage approach in which conventional machine learning generates initial draft annotations, which are then refined using deep learning.

**Ground truth -** Expert-annotated reference data used as the correct answer for training and evaluating segmentation models.

**Transfer learning -** A technique where a model trained on one dataset/task is adapted to perform well on a different but related dataset/task.

**Volumetric data -** Three-dimensional image data composed of voxels (3D pixels) arranged in a regular grid.

**Probability map -** A grayscale image where each voxel’s intensity (0-1) represents the model’s confidence that the voxel belongs to the target structure.

**Inference -** The process of applying a trained model to new data to generate predictions or segmentations.

**Patch size -** The three-dimensional dimensions (in voxels) of image chunks processed by the neural network during training and inference.

**Epoch -** One complete pass through the entire training dataset during neural network training, after which model weights are updated based on cumulative learning.

**Fine-tuning -** The process of adapting a pre-trained neural network to a new task by continuing training on task-specific data.

## Notes

https://github.com/alexmaltsev/DeepBranchAI

### Competing Interest Statement

The authors have declared no competing interest.

### Summary of Updates

Introduction was expanded to clarify the importance of false positives vs false negatives. Methods were additionally clarified with regard to updated code and demonstration.

